# Oocyte surface proteins EGG-1 and EGG-2 are required for eggshell integrity in *Caenorhabditis elegans*

**DOI:** 10.1101/2025.09.12.675948

**Authors:** Ji Kent Kwah, Shannon Pfeiffer, Mst Gitika Khanom, Aimee Jaramillo-Lambert

## Abstract

Metazoan eggs are surrounded by a specialized coat of extracellular matrix that mediates sperm-egg interactions. This coat is rapidly remodeled after fertilization to form a barrier that prevents polyspermy, protects against environmental insults, and provides structural support to the developing embryo. In *C. elegans* several oocyte surface proteins have been identified that mediate these events. However, whether two of these proteins, EGG-1 and EGG-2, are required for fertilization or downstream events has been unclear. Here, we address this question using more recent advances in genome editing tools through the creation of *egg-1 egg-2* deletions of the endogenous loci. We found that *egg-1 egg-2* oocytes are fertilization competent and form rudimentary eggshells. While the integrity of the *egg-1 egg-2* eggshells are compromised and often rupture within the uterus, surprisingly, some embryos are capable of undergoing several rounds of cell division. Overall, our findings demonstrate that EGG-1 and EGG-2 are not required for fertilization but are involved in post-fertilization processes.

## Introduction

The surface of metazoan eggs is covered by a specialized extracellular matrix (ECM) that aids in species-specific egg-sperm interactions. Following fertilization, the ECM is rapidly remodeled to prevent additional sperm entry and to protect the developing embryo (Wong and Wessel. 2006). ECM remodeling occurs as part of egg activation. Egg activation is a critical process that transforms a developmentally quiescent, fertilization-competent oocyte into a fertilization-incompetent, developmentally active one-cell embryo.

In *C. elegans*, egg activation occurs immediately after fertilization and triggers a multitude of changes, including the resumption of meiosis, cortical granule exocytosis, and remodeling of the oocyte ECM. A major outcome of ECM remodeling is eggshell formation, which is essential for embryonic development. The eggshell serves multiple roles, including preventing polyspermy (when more than one sperm fertilizes a single oocyte), regulating the osmotic and chemical environment, and providing structural support to the developing embryo (Stein and Golden. 2018). The *C. elegans* eggshell is composed of multiple layers, assembled in a hierarchical fashion starting with the outermost layer. The outermost three layers– the vitelline layer, chitin layer, and chondroitin proteoglycan (CPG) layer– form the trilaminar outer eggshell. These are followed by the extra-embryonic matrix, the permeability barrier, and finally the peri-embryonic space. Proper coordination between egg activation and the proteins involved in eggshell formation is essential to ensure successful embryogenesis (Olson *et al*. 2012; Stein and Golden. 2018).

Among the proteins found on the oocyte surface, the role of EGG-1 and EGG-2 remains somewhat enigmatic. These two paralogous proteins share 67% amino acid identity and are in close proximity on chromosome III. EGG-1/2 are predicted to be LDL receptor repeat-containing proteins (Kadandale *et al*. 2005). EGG-1/2 localize to the oocyte plasma membrane and are endocytosed after fertilization (González *et al*. 2018; Kadandale *et al*. 2005). However, their precise roles remain unclear due to conflicting experimental results. Prior studies have reported that *egg-1/2* deficient oocytes are fertilization incompetent (Kadandale *et al*. 2005; Lee and Schedl. 2001; Maeda *et al*. 2001). In contrast, Johnston et al. (2010) found that *egg-1/2* deficient oocytes are fertilization-competent but exhibit polyspermy and defects in chitin layer formation. These discrepancies may stem from technical limitations, as earlier studies relied on RNAi-mediated depletion, which can have off-target effects or inefficient mRNA knockdown. In this report, we used CRISPR/Cas9 to generate an *egg-1 egg-2* double-knockout mutant. Our findings indicate that these oocytes are fertilization competent but exhibit significant eggshell defects, including polyspermy, disrupted chitin layer formation, and an impaired permeability barrier.

## Materials and Methods

### *C. elegans* Strains

Standard culturing conditions were used to maintain the *C. elegans* strains used in this study (Brenner. 1973). All worms were maintained on Modified Youngren’s, Only Bacto-peptone (MYOB) plates seeded with *E. coli* OP50. The following strains were utilized: N2, AJL148: *egg-1(ude56) egg-2(ude52)/qC1* III (this study), AJL173: *egg-1(ude56) egg-2(ude52)/qC1* III; *fog-2(oz40)* V (this study), and DG4915: *his-72(uge30[gfp::his-72])* III; *fog-2(oz40)* V.

### CRISPR/Cas-9 mediated genome editing

CRISPR/Cas-9-mediated genome deletions were conducted using the clone-free homology directed repair method with *dpy-10* as a co-CRISPR marker (Arribere *et al*. 2014; Paix *et al*. 2015). Injections were conducted using an injection mix of 1.53 µM Cas9 protein (IDT), 6.4 µM universal tracrRNA (IDT), 1.25 µM *dpy-10* crRNA, 5 µM allele specific crRNA, 0.92 µM *dpy-10* repair oligonucleotide and 2.2 µM allele specific oligonucleotide. All crRNA and oligonucleotide sequences are listed in **Supplementary Table 1**. Genome editing was achieved by injecting each injection mix into the gonad of young adult *+/qC1* hermaphrodites. The F1 generation was screened for edits through PCR. Edited strains were verified by Sanger sequencing. Primers used for PCR and sequencing are as follows: *egg-1* forward TCGCCCAACCCTAACTTGAT, *egg-1* internal reverse TCATCCAACCTTTGCAGCAC, *egg-1* reverse CTTCGGATGTGCTGATCTGC, *egg-2* forward TACTGGTTATTTCGGCGGGA, *egg-2* internal reverse GCTGATCCATGCGATGACTG, *egg-2* reverse TTTGAACAATTCCCCTCGCG.

### Embryonic viability assay and brood sizing

Embryonic viability and brood size assays were conducted using protocols described in (Kwah and Jaramillo-Lambert. 2023). Individual L4 hermaphrodites were transferred onto a single 35 mm MYOB plate spotted with *E. coli* OP50. Each hermaphrodite was allowed to lay embryos at 20°C for 24 hours before being transferred to a new spotted 35 mm MYOB plate until cessation of embryo production. Each plate was screened for hatched larvae and unhatched embryos after 48 hours. Percent embryonic viability was calculated by dividing the total hatched larvae with the total brood size (hatched larvae plus unhatched embryos).

### Widefield Microscopy

The images in Figures 2A, 3, and 4 were acquired with an AxioObserver inverted widefield microscope (Carl Zeiss Inc. Gottingen, Germany) using a 20X Plan-Neofluar (numerical aperture 0.5) or a 40X Plan-Neofluar (numerical aperture 1.3) objective lens and an Axiocam 503 camera (Carl Zeiss Inc.). Each image is of a single focal plane.

Image acquisition, processing and analysis were conducted via Zen Microscopy software (Carl Zeiss Inc. Gottingen, Germany) and ImageJ (Fiji) (Schindelin *et al*. 2012). All images obtained for each respective experiment were obtained using identical parameters, with brightness and contrast adjusted for better visualization.

### Confocal microscopy

The images in Figure 2B were acquired with an Andor Dragonfly spinning disc confocal microscope (Oxford Instruments) using a Plan Apo 63X objective lens (numerical aperture 1.47) and a Zyla sCMOS camera (Oxford Instruments). Image processing and analysis were conducted using Imaris image analysis software (Oxford Instruments). All images were obtained using identical parameters, with brightness and contrast adjusted for better visualization.

### *In utero* imaging of fertilized eggs

*egg-1(ude56) egg-2(ude52)/qC1*; *fog-2(oz40)* (control) or *egg-1(ude56) egg-2(ude52)*; *fog-2(oz40)* females were staged by picking L4 larvae and allowing them to grow into young adults overnight. *gfp::his-72; fog-2(oz40)* adult males were placed with the control or mutant female young adults and allowed to mate for one hour. After mating, the females were immobilized on a 2% agarose pad with 20 µl of 2 mM tetramisole. Imaging of live animals was conducted on an Andor Dragonfly spinning disc confocal microscope (Oxford Instruments) or on an AxioObserver inverted widefield microscope (Carl Zeiss Inc. Gottingen, Germany). For images captured with the AxioObserver widefield microscope, each sample was imaged in a single focal plane. Imaging conducted with the Andor Dragonfly spinning disc confocal microscope were Z-stack projections with 0.5 μm steps for a total of 20 μm.

### Eggshell staining and permeability assays

L4 hermaphrodites of the indicated genotypes were plated onto fresh MYOB plates and allowed to grow for 24 hours at 20°C. Embryos were dissected from the indicated strains in egg buffer [4 mM HEPES (pH 7.4), 94 mM NaCl, 3.2 mM KCl, 2.7 mM CaCl_2_, and 2.7 mM MgCl_2_] supplemented with 16 mM of FM4-64 (Invitrogen) (Bai *et al*. 2020) or Calcofluor White Stain (Sigma-Aldrich) at a ratio of 1:5 Calcofluor White Stain to egg buffer. Dissection of hermaphrodites was conducted with 5 µl of supplemented egg buffer on a cover slip. A depression microscopy slide, to prevent pressure on the eggs, was fixed to the cover slip using four drops of Vaseline on the edges.

### Spermatheca imaging

To image sperm in the spermatheca, L4 hermaphrodites were staged on MYOB plates and were prepared for imaging at three different time points: 16 hours, 24 hours, and 48 hours post-L4. For whole mount DAPI staining, 5-15 hermaphrodites were picked into 5 μL of M9 buffer on a slide. After removal of the M9 with a Kimwipe, worms were fixed by adding 12 μL of room temperature 100% methanol. 15 μL of 2 μg/mL DAPI was added immediately after the methanol evaporated and the samples covered with a coverslip. Slides were incubated in the dark at room temperature for a minimum of 20 minutes before imaging.

### Imaging oocytes in proximal gonad

L4 hermaphrodites were staged on MYOB plates and were prepared for imaging at three different time points: 16 hours, 24 hours, and 48 hours post-L4. Adult hermaphrodites were immobilized on a 2% agarose pad with 20 µl of 2 mM tetramisole. For each sample, images were obtained with DIC optics in a single focal plane.

### Statistical analysis

Statistical analyses were performed using GraphPad Prism 6. Non-parametric Student’s T-test analysis was utilized to determine the statistical significance for the embryonic viability and brood size assays. All experiments were conducted with a minimum of three biological replicates. Error bars indicated standard deviation.

### Data availability

All strains generated in this study are available upon request. The authors state that all data necessary for confirming the conclusions are presented in the article and figures. Supplementary table 1 contains a list of the sequences used for genome editing.

Supplemental material is available online at Genetics.

## Results

### *egg-1* and *egg-2* null mutants are infertile

To initiate the study of the role of EGG-1 and EGG-2 in reproduction, CRISPR/Cas9 genome editing was used to delete the entire open reading frame of both *egg-1* and *egg-2*. As prior work suggested that the double mutant is either fertilization incompetent or developmentally defective (Johnston *et al*. 2010; Kadandale *et al*. 2005), a *+/qC1* balancer strain was injected to generate the strain *egg-1(ude56) egg-2(ude52)/qC1* that can be maintained by picking heterozygous hermaphrodites (**Fig. 1A**). To investigate the function of EGG-1 and EGG-2 in fertilization and embryonic development, brood sizes and embryonic viability were assessed. Heterozygous *egg-1(ude56) egg-2(ude52)/qC1* hermaphrodites produced healthy populations of progeny (average brood size 251.5 ± 79.1, **Fig. 1B**) of viable embryos (99.0% viable, **Fig. 1C**). In contrast, homozygous *egg-1(ude56) egg-2(ude52)* null mutant hermaphrodites did not produce any viable progeny (**Fig. 1B-C**). These results reconfirm that *egg-1* and *egg-2* play a role in *C. elegans* fertility.

**Figure 1.**
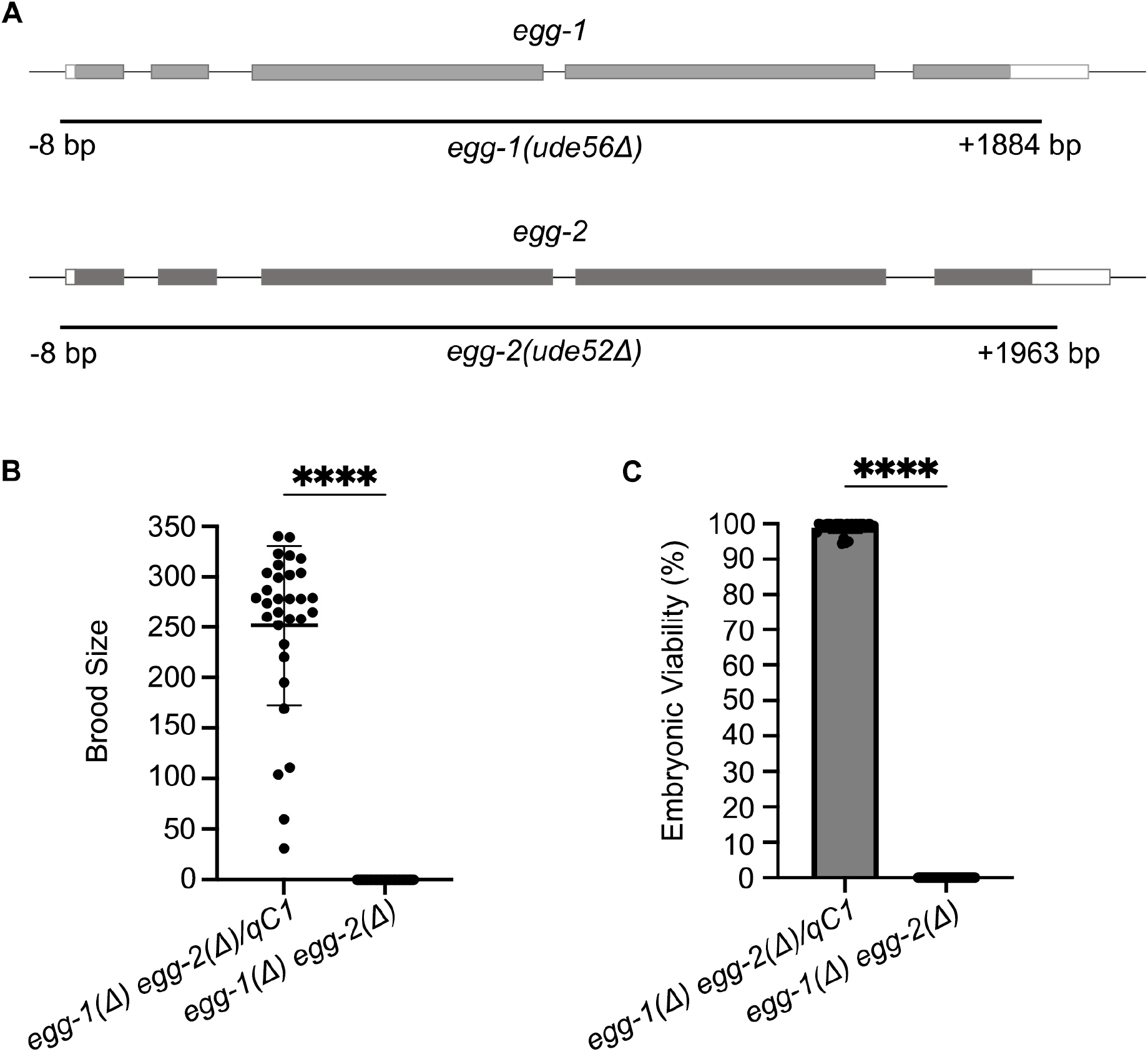
*egg-1 egg-2* null mutants do not produce viable progeny. (A) Representative schematic of *egg-1* and *egg-2* gene structure. Black lines under the gene structures indicate the extent of the CRISPR/Cas9 generated deletions. (B) Brood size counts for control *egg-1(ude56) egg-2(ude52)/qC1* and homozygous *egg-1(ude56) egg-2(ude52)* hermaphrodites. The brood of individual hermaphrodites were plotted from three replicate experiments. The mean brood size is indicated by the black bar. (C) The percentage of viable embryos per brood was determined for both control *egg-1(ude56) egg-2(ude52)/qC1* and homozygous *egg-1(ude56) egg-2(ude52)* mutants. P-values for B & C were calculated by Student’s T-test ****, P<0.0001. *egg-1(ude56) egg-2(ude52)/qC1* N=31. *egg-1(ude56) egg-2(ude52)* N=35.

### *egg-1 egg-2* null mutants are fertilization competent

To evaluate the fertilization competency of *egg-1(ude56) egg-2(ude52)* null mutants, we initially attempted to mate the *egg-1(ude56) egg-2(ude52)* hermaphrodites with males expressing a GFP tagged histone H3.3 (*gfp::his-72*) to mark paternal chromatin. However, this approach was not possible because the sperm contributed by the males were unable to migrate through the uterus of the *egg-1(ude56) egg-2(ude52)* hermaphrodites. While heterozygous hermaphrodites have a uterus filled with embryos fertilized by self-sperm, *egg-1(ude56) egg-2(ude52)* hermaphrodites had uteri filled with ooplasm as the oocytes ruptured during spermathecal transit. We believe this created an environment in which the male sperm were unable to traverse the uterus to reach the spermatheca. Therefore, we generated a feminized strain of *egg-1(ude56) egg-2(ude52)* null mutants by incorporating the *fog-2(oz40)* allele. Ovulation in *C. elegans* is triggered by a sperm-derived hormone-like signal (Miller *et al*. 2001). As *fog-2(oz40)* XX animals do not produce sperm, ovulations will not occur unless male sperm are provided. This ensured spermatids were able to traverse the uterus and enter the spermatheca of the *egg-1(ude56) egg-2(ude52)* mutants before the initiation of ovulation. *In utero* live imaging showed that oocytes of both *egg-1(ude56) egg-2(ude52)/qC1*; *fog-2(oz40)* heterozygous and homozygous *egg-1(ude56) egg-2(ude52)*; *fog-2(oz40)* mutants can be fertilized by male-derived sperm. GFP-labeled paternal chromatin was observed in the most recently formed one-cell embryo passing through the spermatheca, indicating successful fertilization (arrowheads, **Fig. 2A**). To confirm that the sperm was within the newly fertilized egg and not on the outside of the egg, three-dimensional Z-stack live *in utero* spinning disc confocal microscopy was performed. Z-stack projections confirmed the presence of the paternal GFP::H3.3 labeled sperm within newly fertilized eggs (arrowheads, **Fig. 2B**). In addition, we observed some *egg-1(ude56) egg-2(ude52)*; *fog-2(oz40)* embryos had been fertilized by multiple sperm (**Fig. 2C**). From these data we conclude that EGG-1 and EGG-2 are not required for fertilization and may have defects in the block to polyspermy.

**Figure 2.**
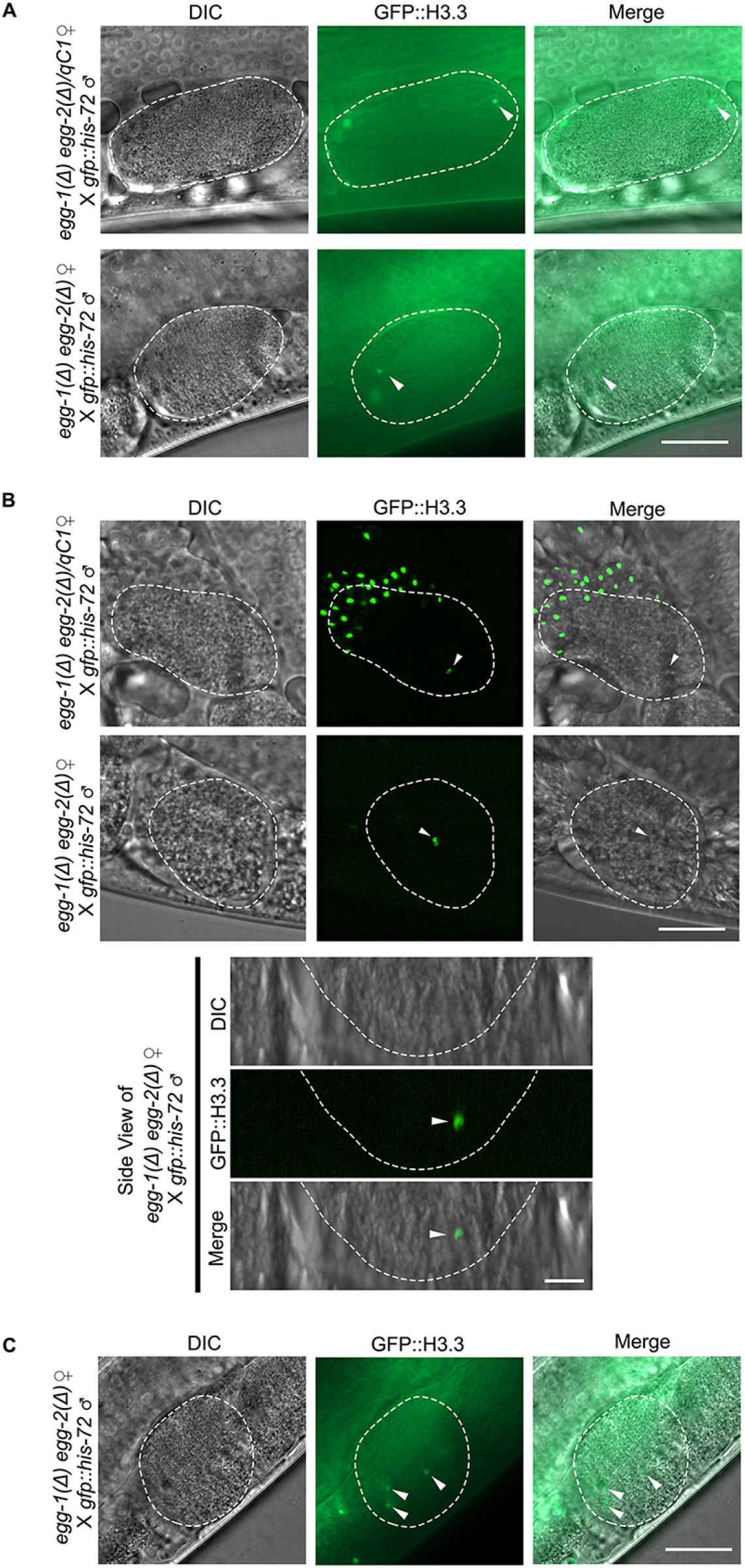
*egg-1 egg-2* null mutants are fertilization competent. (A) Widefield imaging and (B) spinning disc confocal microscopy of feminized control *egg-1(ude56) egg-2(ude52)/qC1; fog-2(oz40)* and homozygous *egg-1(ude56) egg-2(ude52)/qC1; fog-2(oz40)* females mated with males expressing GFP::H3. Green sperm chromatin observed within the newly fertilized embryo indicates the successful fertilization of the *egg-1 egg-2* null oocyte. (C) An example of an embryo with multiple GFP::H3.3-labeled sperm chromatin foci (arrowheads). The white dashed lines outline the fertilized embryos. The total number of fertilized embryos observed per strain between the two imaging techniques: *egg-1(ude56) egg-2(ude52)/qC1; fog-2(oz40)* N=29 and *egg-1(ude56) egg-2(ude52); fog-2(oz40)* N=26. Scale bar = 20 µm.

### EGG-1 and EGG-2 are required for eggshell integrity

During the fertilization assays, we observed that embryos of *egg-1(ude56) egg-2(ude52)* mutants were often ruptured in the uterus of the animal, indicating a defect in eggshell integrity. To further investigate if EGG-1 and EGG-2 play a role in eggshell formation similar to proteins required for egg activation, we assayed for eggshell permeability. Embryos from *egg-1(ude56) egg-2(ude52)* hermaphrodites readily incorporated the lipophilic dye FM4-64 into their plasma membranes, while this dye is excluded from *egg-1(ude56) egg-2(ude52)/qC1* embryos that have intact eggshells (**Fig. 3A**). The *egg-1(ude56) egg-2(ude52)* embryos were also osmotically sensitive with some embryos rupturing during the imaging process (**Fig. 3A**, middle panels). As chitin deposition is required for the major structural integrity of the embryo and other egg activation proteins are required for proper eggshell formation (González *et al*. 2018; Johnston *et al*. 2010; Maruyama *et al*. 2007; Parry *et al*. 2009; Tsukamoto *et al*. 2025), we next tested if *egg-1(ude56) egg-2(ude52)* embryos form the chitin layer of the eggshell. The eggshell of embryos from heterozygous *egg-1(ude56) egg-2(ude52)/qC1* hermaphrodites stain brightly with Calcofluor white, which stains chitin (**Fig. 3B**).

**Figure 3.**
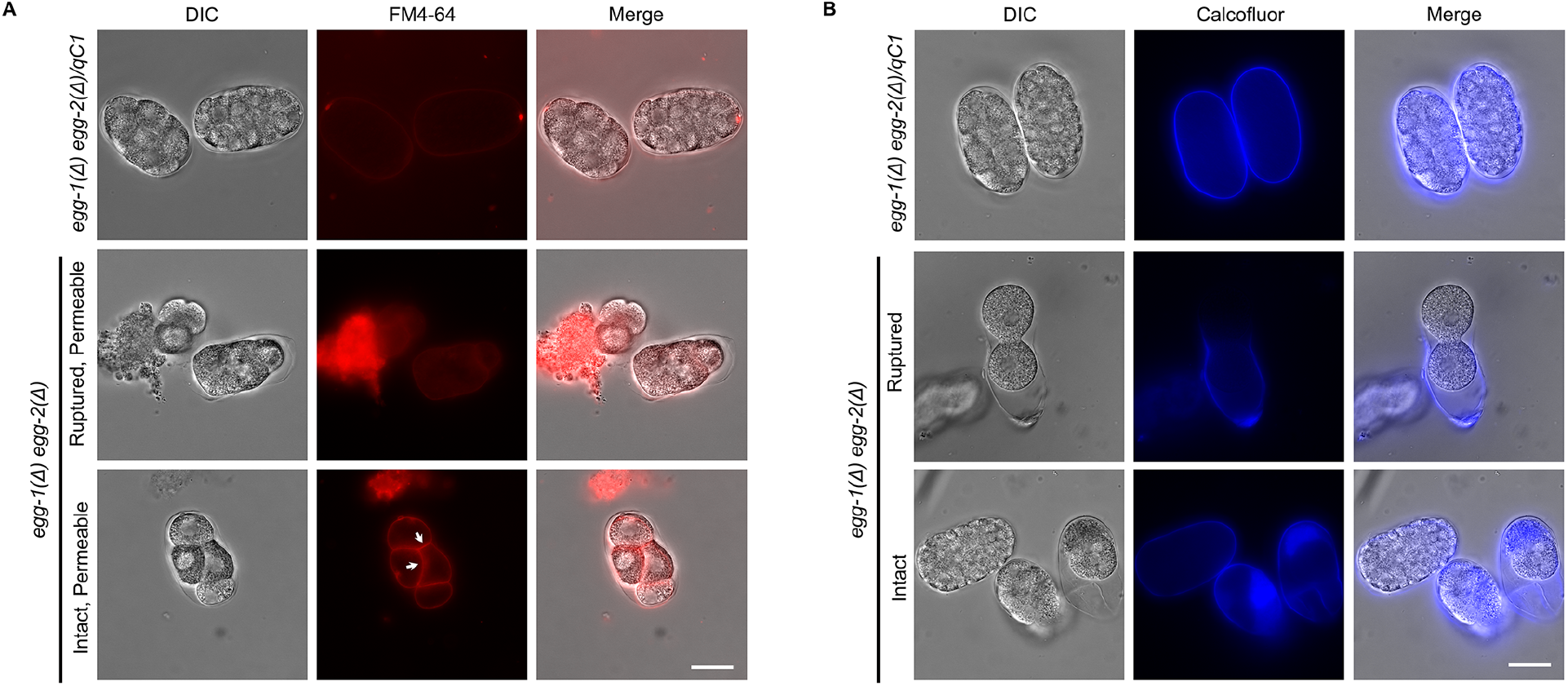
Eggshell integrity is compromised in *egg-1(Δ) egg-2(Δ)* mutants. (A) FM4-64 lipophilic dye is excluded from reaching membranes in the embryo by the eggshell in control *egg-1(ude56) egg-2(ude52)/qC1* (N=115) but permeates the *egg-1(ude56) egg-2(ude52)* mutants, staining the internal membranes of the embryo (white arrows, N=113). Scale bar = 20 µm. (B) Calcofluor white stain to detect the chitin layer of the eggshell. In control *egg-1(ude56) egg-2(ude52)/qC1* a distinct chitin layer (blue) surrounds the developing embryos (N=135). In *egg-1(ude56) egg-2(ude52)* mutant embryos the chitin layer is present but is fragile and prone to rupture (N=111). Scale bar = 20 µm. The scale bar for the side view panels = 10 µm.

Surprisingly, embryos from homozygous *egg-1(ude56) egg-2(ude52)* hermaphrodites also had Calcofluor white staining around the entire eggshell (**Fig. 3B**). This differs from previously published phenotypes of genes involved in egg activation and eggshell integrity (i.e., *cbd-1, spe-11, oops-1*), which only form a chitin cap at the site of sperm entry (Johnston *et al*. 2010; Tsukamoto *et al*. 2025). Ruptured embryos were also observed during these assays, with some eggshells seeming to slough off of the embryo (**Fig. 3B** middle panels). We also observed that some *egg-1(ude56) egg-2(ude52)* embryos completed several rounds of cell division without rupturing (**Fig. 3A-B**). This phenotype differs from egg activation mutants, which fail to undergo cell division (Hill *et al*. 1989; Johnston *et al*. 2010; Maruyama *et al*. 2007; Tsukamoto *et al*. 2025).

### *egg-1 egg-2* null mutants deplete sperm more rapidly than controls

Despite the fertility and eggshell defects observed in the *egg-1(ude56) egg-2(ude52)* hermaphrodites, these worms did not have any obvious gametogenesis or somatic defects. However, we observed that oocytes in the *egg-1(ude56) egg-2(ude52)* gonad appeared to have a stacking phenotype in older adults. To investigate this further, the gonads of young adult hermaphrodites were imaged at 16 h, 24 h, and 48 h post-L4. In wild-type *C. elegans*, oocytes in the most proximal gonad are large with a cuboidal shape. The oocyte closest to the spermatheca will receive the meiotic maturation/ovulation signal from sperm in the spermatheca where it will become rounder in shape as it matures (McCarter *et al*. 1999). Both wild-type (N2) and *egg-1(ude56) egg-2(ude52)* mutant gonads display normal appearing oocytes at 16 h and 24 h post-L4. However, by 48 h oocytes in *egg-1(ude56) egg-2(ude52)* mutant hermaphrodites stacked up in the gonad arm (**Fig. 4A**). This phenotype is associated with depletion of a sperm-provided meiotic maturation/ovulation signal (McCarter *et al*. 1999; Miller *et al*. 2001). Loss of the sperm signal can occur through several different mechanisms including sperm motility defects or rapid consumption of sperm due to polyspermy. To determine if sperm were depleted more rapidly in *egg-1(ude56) egg-2(ude52)* mutants, we fixed and DAPI stained whole worms and took images of spermathecae. DAPI stained sperm can be found in the spermathecae of day 1 (24 h) and day 2 (48 h), and day 3 wild-type adults (**Fig. 4B**). However, in *egg-1(ude56) egg-2(ude52)* mutants very few sperm were observed in the spermathecae of day 2 and day 3 adults (**Fig. 4B**). We believe this rapid depletion of sperm may be due, in part, to polyspermy as we observed instances of polyspermy when performing the fertilization assays (**Fig. 2C**). However, this will need to be investigated further in the future.

**Figure 4.**
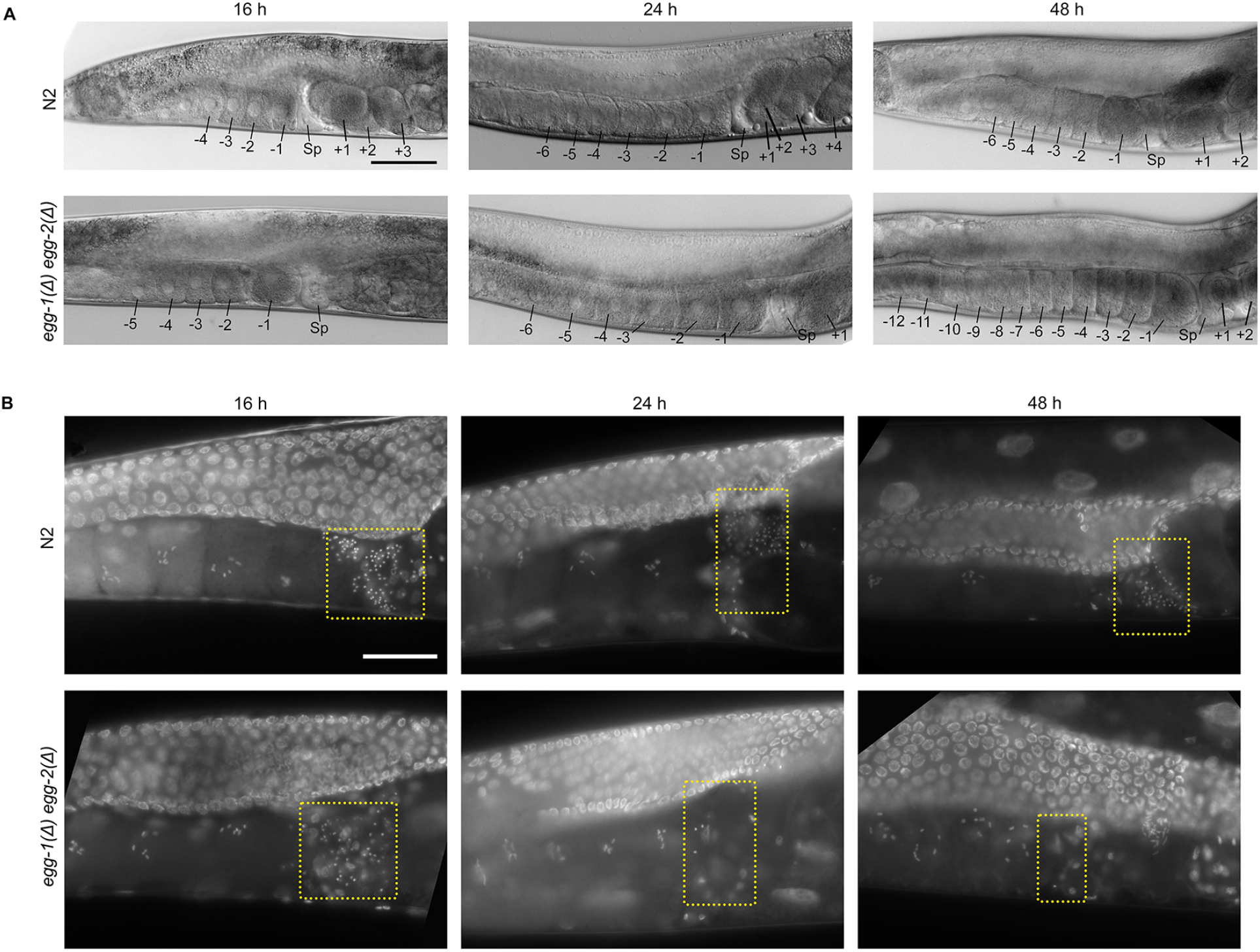
Oocyte stacking and sperm depletion in *egg-1(Δ) egg-2(Δ)* mutants. (A) DIC images of the proximal gonads of wild-type (N2) and *egg-1(ude56) egg-2(ude52)* hermaphrodites. N2 N=83; *egg-1(ude56) egg-2(ude52)* N=61. Proximal oocytes are labeled with negative numbers (e.g., -1 to -4 in N2 at 16 h), newly fertilized embryos are indicated with positive numbers (e.g., +1 to +3 in N2 at 16 h), and Sp marks the spermatheca. By 48 h post-L4, *egg-1(ude56) egg-2(ude52)* mutant hermaphrodites have many more oocytes in the proximal gonad (>6) indicating a “stacking” phenotype. Scale bar = 50 µm. (B) Images of the proximal region of DAPI stained hermaphrodite gonads. Yellow dashed boxes indicate the region of the spermatheca where sperm are stored. Sperm is depleted more rapidly from *egg-1(ude56) egg-2(ude52)* mutants than from wild-type (N2) hermaphrodites. By 48 h post-L4, only a few spermatids are visible in the *egg-1(ude56) egg-2(ude52)* spermathecae. Scale bar = 20 µm. N2 N=96; *egg-1 egg-2* N=89.

## Discussion

In this study, we utilized more recent genome editing techniques to create an *egg-1 egg-2* null mutant. We found that the oocyte plasma membrane proteins EGG-1 and EGG-2 are not required for fertilization competency. This differs from previous findings where knockdown of *egg-1* and *egg-2* via RNAi resulted in a sterile phenotype with unfertilized oocytes in the uterus (Kadandale *et al*. 2005; McCarter *et al*. 1999).

However, in these experiments, RNAi targeting *egg-1* was found to also deplete *egg-2*. As there were off target effects due to high sequence similarity, one possibility is that under these conditions additional, yet undiscovered, *egg* genes were knocked down causing the sterility defects.

Our results support the findings of Johnston et. al., 2010, which demonstrated that simultaneous RNAi knockdown of *egg-1* and *egg-2* in hermaphrodites resulted in fertilized embryos with eggshell defects (**Fig. 2-3**). However, we did observe phenotypic differences from this previous study. The Johnston et. al., 2010 study found that *egg-1(RNAi) egg-2(RNAi)* animals produce severely fragmented fertilized eggs with fragmented eggshells. In this study, we found that some *egg-1(ude56) egg-2(ude52)* embryos underwent several rounds of cell division (**Fig. 3A-B**). We also found that *egg-1(ude56) egg-2(ude52)* embryos formed a continuous albeit permeable eggshell (**Fig. 3A-B**).

Current models place EGG-1 and EGG-2 on the surface of the oocyte plasma membrane where it either interacts with as yet unknown sperm surface proteins or with CBD-1 (chitin binding domain protein) and the egg activation complex (EGG-3, CHS-1, EGG-4/5, and MBK-2). Our data suggests that EGG-1 and EGG-2 are not involved in fertilization; however, the results of this study cannot definitively place them in the egg activation pathway. The *egg-1(ude56) egg-2(ude52)* phenotypes we observed are less severe than *cbd-1* mutants-CBD-1 regulates eggshell assembly and anchors egg activation complex proteins- and egg activation mutants. *cbd-1* and egg activation mutants do not divide and have severely disrupted eggshells, whereas all *egg-1(ude56) egg-2(ude52)* embryos make rudimentary eggshells and some undergo cell division (González *et al*. 2018; Hill *et al*. 1989; Johnston *et al*. 2010; Parry *et al*. 2009; Tsukamoto *et al*. 2025; Zhang *et al*. 2005). We also observed some examples of polyspermy (**Fig. 2C**), a feature of some egg activation mutants; however, this needs to be investigated further to determine the extent of the defect .

There is evidence that EGG-1 and EGG-2 interact with both CBD-1 and egg activation proteins. EGG-1 was not localized on the oocyte plasma membrane when *cbd-1* was knocked down (RNAi) and when *egg-1* and *egg-2* are co-depleted by RNAi both CBD-1 and CHS-1 (chitin synthase and part of egg activation complex) have an uneven distribution on the plasma membrane (González *et al*. 2018; Johnston *et al*. 2010). Direct protein-protein interactions have not been determined between these various components. Future studies to determine both the genetic hierarchy, molecular activity, and protein interactions of EGG-1 and EGG-2 with the various eggshell and egg activation proteins will provide crucial knowledge on how animal cells remodel egg coats after fertilization and support transition from quiescence to active development.

## Acknowledgments

We are grateful to David Greenstein for sharing the feminized GFP::H3.3 strain. We thank Wormbase as an invaluable tool for the *C. elegans* research community. We are especially thankful to members of the Jaramillo-Lambert laboratory for helpful discussions and critical feedback of this manuscript.

## Author Contributions

Design: JKK and AJL. Experiments: JKK, SP, and GK. Analysis: JKK, SP, GK, and AJL. Writing: JKK, SP, and AJL.

## Literature Cited

Arribere, J. A., R. T. Bell, B. X. H. Fu, K. L. Artiles, P. S. Hartman et al, 2014 Efficient marker-free recovery of custom genetic modifications with CRISPR/Cas9 in Caenorhabditis elegans. Genetics 198: 837–846.

Bai, X., L. Huang, S. Chen, B. Nebenfuehr, B. Wysolmerski et al, 2020 Loss of the seipin gene perturbs eggshell formation in Caenorhabditis elegans. Development 147: dev192997. doi: 10.1242/dev.192997.

Brenner, S., 1973 The genetics of behaviour. Br. Med. Bull. 29: 269–271.

González, D. P., H. V. Lamb, D. Partida, Z. T. Wilson, M. Harrison et al, 2018 CBD-1 organizes two independent complexes required for eggshell vitelline layer formation and egg activation in C. elegans. Dev. Biol. 442: 288–300.

Hill, D. P., D. C. Shakes, S. Ward and S. Strome, 1989 A sperm-supplied product essential for initiation of normal embryogenesis in Caenorhabditis elegans is encoded by the paternal-effect embryonic-lethal gene, spe-11. Dev. Biol. 136: 154–166.

Johnston, W. L., A. Krizus and J. W. Dennis, 2010 Eggshell chitin and chitin-interacting proteins prevent polyspermy in C. elegans. Current Biology 20: 1932.

Kadandale, P., A. Stewart-Michaelis, S. Gordon, J. Rubin, R. Klancer et al, 2005 The egg surface LDL receptor repeat-containing proteins EGG-1 and EGG-2 are required for fertilization in Caenorhabditis elegans. Curr. Biol. 15: 2222–2229.

Kwah, J. K., and A. Jaramillo-Lambert, 2023 Measuring embryonic viability and brood size in Caenorhabditis elegans. J. Vis. Exp. (192):10.3791/65064. doi: 10.3791/65064.

Lee, M. H., and T. Schedl, 2001 Identification of in vivo mRNA targets of GLD-1, a maxi-KH motif containing protein required for C. elegans germ cell development. Genes Dev. 15: 2408–2420.

Maeda, I., Y. Kohara, M. Yamamoto and A. Sugimoto, 2001 Large-scale analysis of gene function in Caenorhabditis elegans by high-throughput RNAi. Curr. Biol. 11: 171– 176.

Maruyama, R., N. V. Velarde, R. Klancer, S. Gordon, P. Kadandale et al, 2007 EGG-3 regulates cell-surface and cortex rearrangements during egg activation in Caenorhabditis elegans. Current Biology 17: 1555.

McCarter, J., B. Bartlett, T. Dang and T. Schedl, 1999 On the control of oocyte meiotic maturation and ovulation in Caenorhabditis elegans. Dev. Biol. 205: 111–128.

Miller, M. A., V. Q. Nguyen, M. H. Lee, M. Kosinski, T. Schedl et al, 2001 A sperm cytoskeletal protein that signals oocyte meiotic maturation and ovulation. Science 291: 2144–2147.

Olson, S. K., G. Greenan, A. Desai, T. Muller-Reichert and K. Oegema, 2012 Hierarchical assembly of the eggshell and permeability barrier in C. elegans. J. Cell Biol. 198: 731–748.

Paix, A., A. Folkmann, D. Rasoloson and G. Seydoux, 2015 High efficiency, homology-directed genome editing in Caenorhabditis elegans using CRISPR-Cas9 ribonucleoprotein complexes. Genetics 201: 47–54.

Parry, J. M., N. V. Velarde, A. J. Lefkovith, M. H. Zegarek, J. S. Hang et al, 2009 EGG-4 and EGG-5 link events of the oocyte-to-embryo transition with meiotic progression in C. elegans. Curr. Biol. 19: 1752–1757.

Schindelin, J., I. Arganda-Carreras, E. Frise, V. Kaynig, M. Longair et al, 2012 Fiji: An open-source platform for biological-image analysis. Nat. Methods 9: 676–682.

Stein, K. K., and A. Golden, 2018 The C. elegans eggshell. WormBook 2018: 1–36.

Tsukamoto, T., J. K. Kwah, M. E. Zweifel, N. Courtemanche, M. D. Gearhart et al, 2025 A sperm-oocyte protein partnership required for egg activation in Caenorhabditis elegans. Development 152: dev204674. doi: 10.1242/dev.204674. Epub 2025 Jun 25.

Wong, J. L., and G. M. Wessel, 2006 Defending the zygote: Search for the ancestral animal block to polyspermy. Curr. Top. Dev. Biol. 72: 1–151.

Zhang, Y., J. M. Foster, L. S. Nelson, D. Ma and C. K. S. Carlow, 2005 The chitin synthase genes chs-1 and chs-2 are essential for C. elegans development and responsible for chitin deposition in the eggshell and pharynx, respectively. Dev. Biol. 285: 330–339.

